# Scaling probabilistic models of genetic variation to millions of humans

**DOI:** 10.1101/013227

**Authors:** Prem Gopalan, Wei Hao, David M. Blei, John D. Storey

## Abstract

One of the major goals of population genetics is to quantitatively understand variation of genetic polymorphisms among individuals. To this end, researchers have developed sophisticated statistical methods to capture the complex population structure that underlies observed genotypes in humans, and such methods have been effective for analyzing modestly sized genomic data sets. However, the number of genotyped humans has grown significantly in recent years, and it is accelerating. In aggregate about 1M individuals have been genotyped to date. Analyzing these data will bring us closer to a nearly complete picture of human genetic variation; but existing methods for population genetics analysis do not scale to data of this size. To solve this problem we developed TeraStructure. TeraStructure is a new algorithm to fit Bayesian models of genetic variation in human populations on tera-sample-sized data sets (10^12^ observed genotypes, e.g., 1M individuals at 1M SNPs). It is a principled approach to Bayesian inference that iterates between subsampling locations of the genome and updating an estimate of the latent population structure of the individuals. On data sets of up to 2K individuals, TeraStructure matches the existing state of the art in terms of both speed and accuracy. On simulated data sets of up to 10K individuals, TeraStructure is twice as fast as existing methods and has higher accuracy in recovering the latent population structure. On genomic data simulated at the tera-sample-size scales, TeraStructure continues to be accurate and is the only method that can complete its analysis.

**Software:** TeraStructure is available for download at https://github.com/premgopalan/terastructure.

**Funding:** This research was supported in part by NIH grant R01 HG006448 and ONR grant N00014-12-1-0764.

## INTRODUCTION

The quantitative characterization of genetic variation in human populations plays a key role in understanding evolution, migration, and trait variation. Genetic variation of humans is highly structured in that frequencies of genetic polymorphisms are heterogeneous among human sub-populations. Therefore, to comprehensively understand human genetic variation, we must also understand the underlying structure of human populations.

Over the last fifteen years, scientists have successfully used genome-wide Bayesian models of genetic polymorphisms to infer the latent structure embedded in an observed population. The probabilistic model of Pritchard, Stephens and Donnelly [1], which we will refer to as the “PSD model”, has become a standard tool both for exploring hypotheses about human genetic variation and accounting for latent population structure in downstream analyses. The basic idea behind the PSD model is that each individual’s ancestry is composed of a mixture of ancestral populations, and an individual’s genotype can thus be modeled as a random process that mixes the frequencies of genetic variants from among these ancestral populations.

The PSD model turns the problem of estimating population structure into one of posterior inference, computing the conditional distribution of hidden random variables given observed random variables. The hidden random variables in the model encode the underlying ancestral structure; the observed random variables are the genotypes among a sample of individuals and SNPs. As for many modern Bayesian models, however, this posterior is not tractable to compute: the original algorithm for using the PSD model [1] and subsequent innovations [2; 3] are all methods for approximating it. It is through these approximations of the posterior that population geneticists can explore the latent population structure of their data and account for ancestry in downstream analyses.

Modern genetics, however, cannot take full advantage of the PSD model and related probabilistic models. The reason is that the existing solutions to the core computational problem—the problem of estimating the latent structure given a collection of observed genetic data—cannot handle the scale of modern datasets. These solutions require repeatedly cycling through the entire data set, and repeatedly refining estimates of the latent population structure. With massive data sets available today, this is not a practical methodology.

Specifically, the sample sizes of genome-wide association studies now routinely involve tens of thousands of people. Furthermore, public and private initiatives have managed to measure genome-wide genetic variation on hundreds of thousands of individuals. For example, a recent study by the company 23andMe collected genome-wide genotypes from 162,721 individuals [4]. Taken together, we now have dense genome-wide genotype data on the order of a million individuals. Fitting probabilistic models on these data would provide an unprecedented characterization of genetic variation and the structure of human populations. But, as we show in our study, this analysis is not possible with the current state of the art.

To solve this problem, we have developed TeraStructure, an algorithm for analyzing data sets of up to 10^12^ genotypes (and even more with advanced computing architectures). It is based on a scalable implementation of “variational inference” [5; 6], a general optimization-based strategy for Bayesian computation. TeraStructure’s computational flow iterates between subsampling observed single nucleotide polymorphism (SNP) genotypes, analyzing the subsample, and updating its estimate of the hidden ancestral populations. This flow enables us to analyze massive data sets with the PSD model and can be adapted to other statistical models as well [7]. It changes the scale at which we can use Bayesian models of population genetics.

## RESULTS

TeraStructure provides a statistical estimate of the PSD model, capturing the heterogenous mixtures of ancestral populations that are inherent in a data set of observed human genomes. Formally, the PSD model assumes that there are *K* ancestral populations, each characterized by its minor allele frequencies *β*_*k*_ for each of the SNPs. Further, it assumes that each individual in the sample exhibits those populations with different proportions *θ*_*i*_. Finally, it assumes that each SNP genotype *l* in each individual *i*, denoted by *x*_*i,l*_, is drawn from an ancestral population that itself is drawn from the individual-specific proportions. If we code each SNP genotype as a 0, 1, or 2 (to denote the three possible genotypes), then it models *x*_*i,l*_ *∼* Binomial(2, *p*_*i,l*_) where *p*_*i,l*_ = ∑_*k*_ *θ*_*i*,*k*_ *β*_*k*,*l*_.

Inference is the central computational problem for the PSD model. Given an data set of observed genotypes {*x*_*i,l*_}, we must estimate the per-individual population proportions *θ*_*i*_ and the statistical signature of each latent population, i.e., the minor allele frequencies, *β*_*k*_. These estimates come from the posterior distribution *p*(*θ, β* | *x*), which is the conditional distribution of the latent structure given the observed data. Existing methods solve this problem by cycling between analyzing all the SNPs of all of the individuals, and updating an estimate of the latent populations. This approach is infeasible for massive data sets.

Figure 1 illustrates the computational flow of TeraStructure, which is intrinsically efficient. At each iteration, it maintains an estimate of the population proportions for each person and the allele frequencies for each population.^1^ It repeatedly iterates between the following steps: (a) sample a SNP from the data, *x*_·,*l*_, the measured genotypes at a single marker in the genome across all people, (b) analyze how the current estimates of the ancestral populations explain the genotypes at that SNP, and (c) update the estimates of the latent structure—both the ancestral allele frequencies and per-individual population proportions.

**Figure 1:**
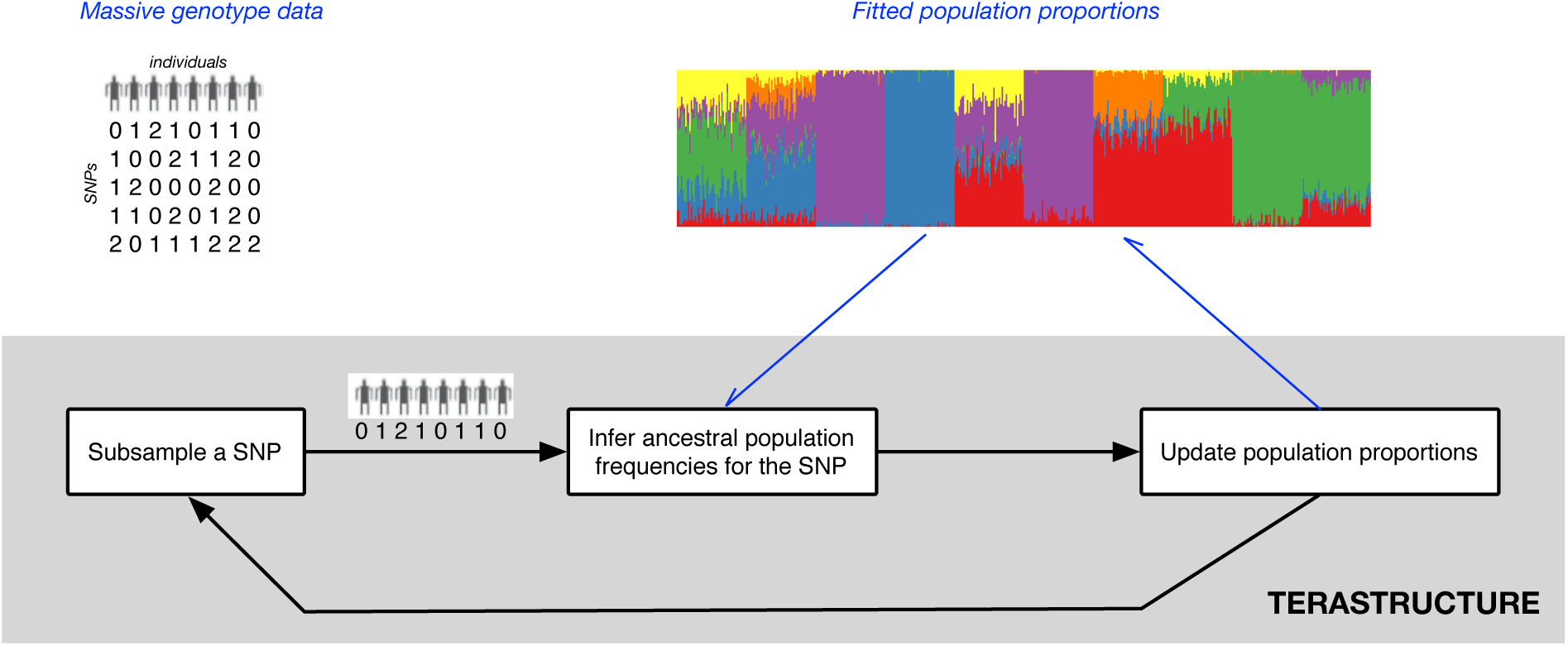
A schematic diagram of TeraStructure, stochastic variational inference for the Pritchard-Stephens-Donnelly (PSD) model. The algorithm maintains an estimate of the latent population proportions for each individual. At each iteration it samples SNP measurements from the large database, infers the per-population frequencies for that SNP, and updates its idea of the population proportions. This is much more efficient than algorithms that must iterate across all SNPs at each iteration.

It is the subsampling step of the inner loop that allows TeraStructure to scale to massive genetic data. Rather than scan the entire population at each iteration, it iteratively subsamples a SNP, analyzes the subsample, and updates its estimate. On small data sets, this leads to faster estimates that are as good as those obtained by the slower procedures. More importantly, it lets us scale the PSD model up to sample sizes that are orders of magnitude larger than what the current state of the art can handle. We further emphasize that the technical approach behind TeraStructure—one that repeatedly subsamples from a massive data set and then updates an estimate of its hidden structure—can be adapted to many Bayesian models that are used in genome analysis, such as HMMs, phylogenetic trees, and others.

TeraStructure is built on variational inference, a method for approximate Bayesian computation that comes from statistical machine learning [5]. The main idea behind variational inference for the PSD model is as follows. We first parameterize individual distributions for each latent variable in the model, i.e., a distribution for each set of per-population allele frequencies *q*(*β*_*k*_) and a distribution for each individual’s population proportions *q*(*θ*_*i*_). We then fit these distributions so that their product is close to the true posterior, where closeness is measured by Kullback-Leibler divergence,

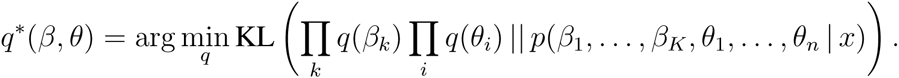

(Kullback-Leibler is an information-theoretic quantity that asymmetrically measures the distance between two distributions.) Thus we do Bayesian inference by solving an optimization problem. The key idea in TeraStructure is to solve this optimization problem with stochastic variational inference [7], a way of doing variational inference that scales to large data. Specifically, we optimize the KL divergence by following noisy realizations of its derivatives, where the noise comes from subsampling the data at each iteration (see Figure 1). A noisy derivative computed from a subsample is much cheaper to compute than the true derivative, which requires iterating over the entire data set. See Methods for the mathematical details that outline the variational objective function and how to compute noisy derivatives from subsamples.

We applied TeraStructure to both real and simulated data sets to study and demonstrate its good performance. We compared it to ADMIXTURE [2] and fastSTRUCTURE [3], the two algorithms for estimating the PSD model that work on modestly sized data. In our comparisons, we timed all the algorithms under equivalent computational conditions. On simulated data, where the truth is known, we measured the quality of the resulting fits by computing the KL divergence between the estimated models and the truth. On the real data sets, where the truth is not known, we measured model fitness by predictive log likelihood of held-out measurements (Methods). The smaller the KL divergence and the larger the predictive likelihood, the better a method performs.

We first analyzed two real data sets: the Human Genome Diversity Panel (HGDP) data set [8; 9] and the 1000 Genomes Project (TGP) [10]. After preprocessing, HGDP consisted of 940 individuals at 642,951 SNPs for a total of 604 million observed genotypes and TGP consisted of 1,718 people at 1,854,622 SNPs for a total of 3.2 billion observed genotypes. In previous work, ADMIXTURE and fastSTRUCTURE have been shown to perform reasonably well on data sets of this size [2; 3]. In applying all three algorithms to these data, we found that TeraStructure equalled the predictive log likelihood of held-out measurements (Table S1) and it completed its estimation in a comparable period of time (Table 1).

**Table 1:** The running time of all algorithms on both real and synthetic data. TeraStructure is the only algorithm that can scale beyond *N* = 10, 000 individuals to the synthetic data sets with *N* = 100, 000 individuals and *N* = 1, 000, 000 individuals. *S* is the fraction of SNP locations sub-sampled, with repetition, during training; *L* is the number of SNP locations. The TeraStructure and ADMIXTURE algorithms were run with ten parallel threads, while fastSTRUCTURE, which does not have a threading option, was run with a single thread. Even under the best-case assumption of ten times speedup due to parallel computation, the TeraStructure algorithm is twice as fast as both ADMIXTURE and fastSTRUCTURE algorithms on the data set with *N* = 10, 000 individuals. On the real data sets, TeraStructure is as fast as the other algorithms. In contrast to other methods that iterated multiple times over the entire data set, TeraStructure iterated over the SNP locations at most once on all data sets.

| Data set | $N$ | $L$ | $S$ | Time (hours) | | |
| --- | --- | --- | --- | --- | --- | --- |
|  |  |  |  | TeraStructure | ADMIXTURE | fastSTRUCTURE |
| HGDP | 940 | 644,258 | 0.9 | < 1 | < 1 | 12 |
| TGP | 1718 | 1,854,622 | 0.5 | 3 | 3 | 21 |
| Scenario A (10K) | 10,000 | 1,000,000 | 1.0 | 9 | 28 | 216 |
| Scenario A (100K) | 100,000 | 1,000,000 | 0.7 | 158 | – | – |
| Scenario A (1M) | 1,000,000 | 1,000,000 | 0.5 | 509 | – | – |
| Scenario B | 10,000 | 100,000 | 1.0 | 6.9 | 31 | 140 |
| Scenario B | 10,000 | 1,000,000 | 0.5 | 9.3 | – | – |

We then studied the algorithms on synthetic data. We designed two simulation scenarios (Figures 2–3, Methods). Scenario A draws population proportions *θ*_*i*_ to be symmetric among the ancestral populations; this matches the assumptions of the PSD model. Scenario B draws *θ*_*i*_ such that there is a spatial relationship between the ancestral populations; this diverges from the underlying assumptions of PSD. We used scenario A to demonstrate the scalability of the methods. Data sets from this scenario contained 10,000 individuals, 100,000 individuals, and 1M individuals, each with 1M SNP genotypes per individual and 6 ancestral populations. We used Scenario B to study the accuracy of the methods under a mismatched model, specifically when there are spatial correlations in population structure [11]. With these simulations we varied the number of SNPs; one data set contained 100,000 SNPs and another contained 1M SNPs, each with 10,000 individuals and 10 ancestral populations.

**Figure 2:**
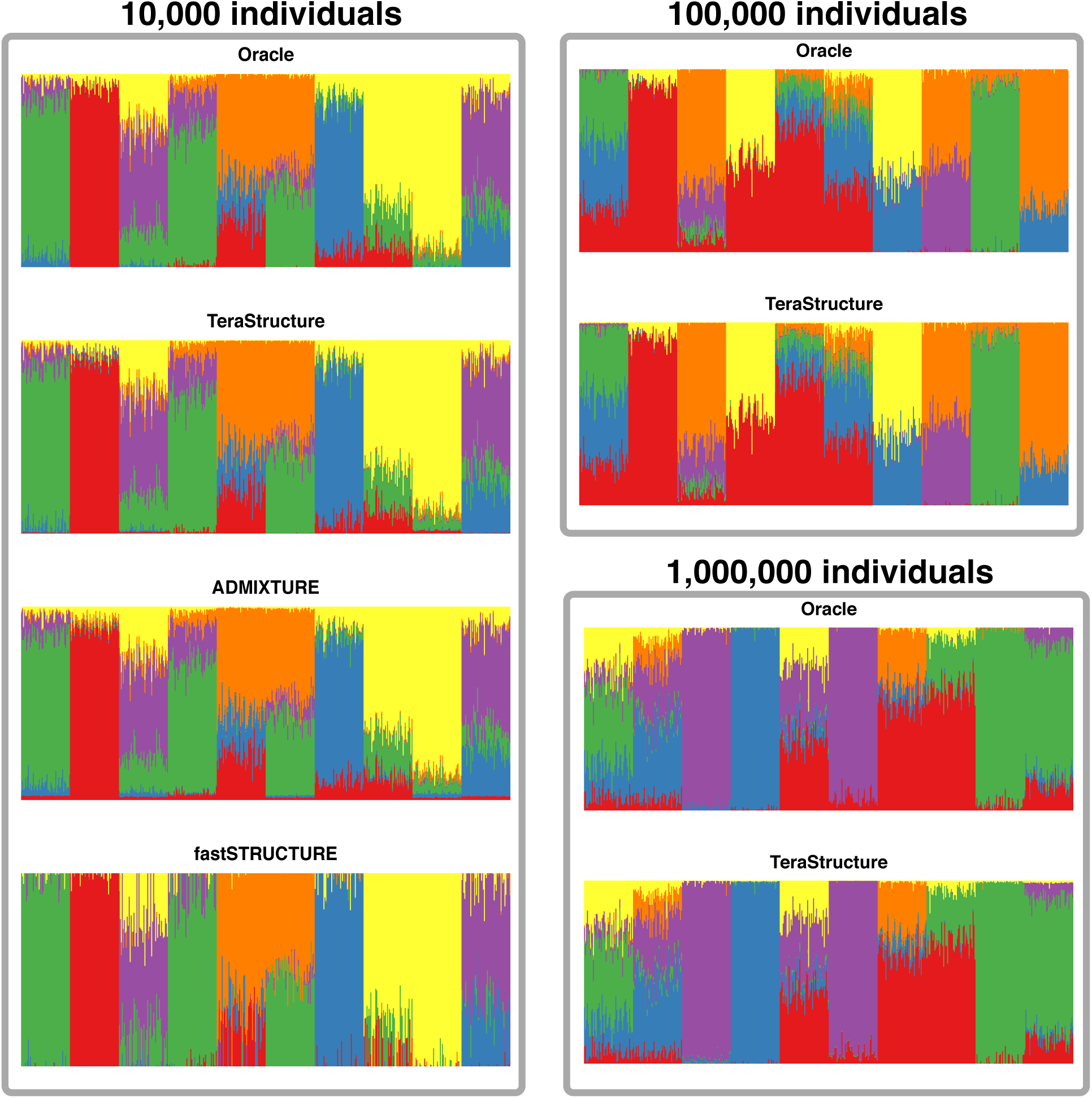
TeraStructure recovers the ground truth per-individual population proportions on the synthetic data sets generated via Scenario A. Each panel shows a visualization of the ground truth 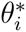 and the inferred 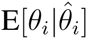 for all individuals in a data set. The current state-of-the-art algorithms cannot complete their analyses of 100,000 and 1,000,000 individuals. TeraStructure is able to analyze data of this size and gives highly accurate estimates.

**Figure 3:**
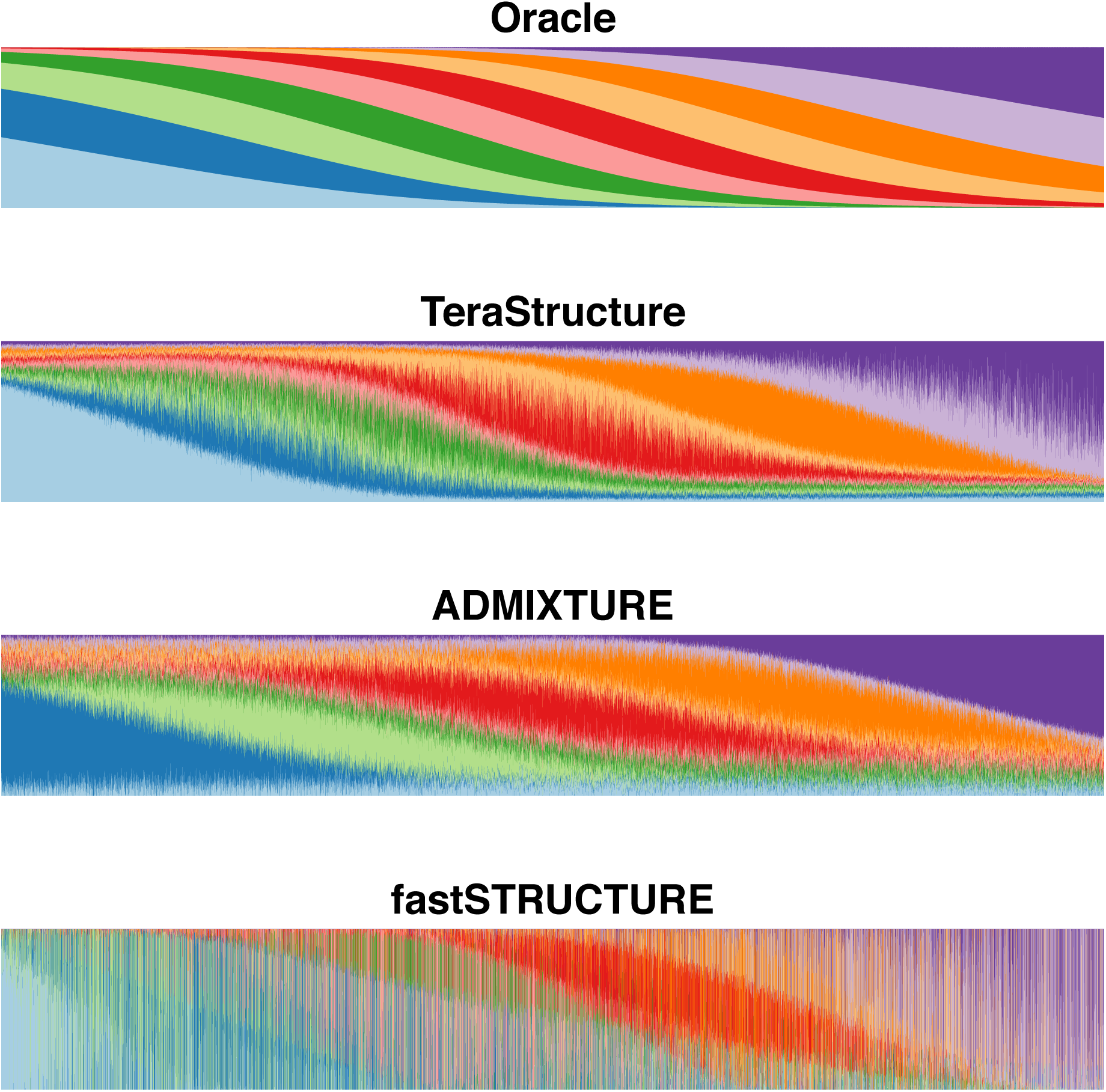
TeraStructure is the most accurate method for Scenario B simulations. The top panel shows the ground truth 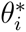, where each individual’s proportions are a function of their position, e.g. the left most individual’s proportions are dominated by the two blue ancestral populations while the right most individual’s proportions are dominated by the two purple ancestral populations. A quantitative measure of the accuracies can be found in Table S2.

On these synthetic data we know the ground truth individual proportions, and we can visualize how well each algorithm reconstructs them (Figures 2–3). For Scenario A, we found that ADMIXTURE and fastSTRUCTURE were only able to analyze the 10,000-individual set, on which TeraStructure was both 2-3 times faster and more accurate (Tables 1 and S2). More importantly, TeraStructure was the only algorithm that was able to analyze the larger data sets of 100,000 individuals and 1M individuals, and again with high accuracy (Figure 2 and Table S2). Further, TeraStructure was by far the fastest and most accurate method on Scenario B (Table S2 and Figure 3).

TeraStructure is an iterative optimization algorithm and thus uses a convergence criterion to decide when to stop iterating (Methods). This lets us gauge how many SNPs were necessary to sample before the algorithm had learned the structure of the population. On the HGDP and TGP data, we found that TeraStructure needed to sample *∼*90% and *∼*50% of the SNPs, respectively, before converging (Table 1). On the tera-sample-sized data set of 1M individuals at 1M SNPs (Scenario A), TeraStructure sampled *∼*50% of the SNPs before converging.

Any analysis with the PSD model requires choosing the number of ancestral populations *K*. TeraStructure addresses this choice using a predictive approach [12]. We hold out a set of genome SNP locations for each individual and compute the average predictive log likelihood under the model for varying numbers of ancestral populations. Our choice of *K* is the one that assigns the highest probability to the held-out set. Our sensitivity analysis revealed that *K* = 8 had the highest validation likelihood on the TGP data, while *K* = 10 had the highest likelihood on the HGDP data (Figure S3). On the real data sets, we fixed the number of populations *K* for each data set to the *K* with the highest validation likelihood (Table S1); on simulated data sets, we set *K* to the number of ground truth ancestral populations (Table 1).

## DISCUSSION

TeraStructure is a scalable algorithm that repeatedly takes strategic subsamples of genotyping data to uncover the underlying structure of human populations. We demonstrated TeraStructure’s superior performance by applying it to large and globally sampled human SNP genotype data, finding equal accuracy and improved speed. Further, we used a comprehensive simulation study to show that TeraStructure can accurately fit a standard probabilistic model of population genetic structure on data sets with a million individuals and 10^12^ observed genotypes. This is orders of magnitude beyond the capabilities of current state-of-the-art algorithms. We note that our results are from a modest computing platform. On advanced computing architectures, TeraStructure can analyze even larger data sets, and holds promise of characterizing the structure of world-scale human populations.

Fitting probabilistic models of population structure, such as the PSD model, is a crucial component of modern population genetics. The size of such studies is growing and thus it is vital that our statistical algorithms scale to millions of individuals and trillions or more genotype observations. We have shown that such analyses are not possible with existing algorithms, which require multiple iterations over the entire data. TeraStructure overcomes this limitation with a more efficient computational flow—one that iterates between subsampling from a population, analyzing the sample, and updating an estimate of hidden structure—without compromising the principles and statistical assumptions behind PSD analysis. Using TeraStructure to analyze tera-sample-size data sets will provide the most comprehensive analyses to date of the global population genetics of humans.

## METHODS

### Data sets

**Real data sets.** We used genotype data from the Human Genome Diversity Project (HGDP) and 1000 Genomes Project (TGP), which are two publicly available datasets that sampled individuals globally. To insure the quality of the data, we filtered individuals for 95% genotyping completeness and we removed SNPs with lower than 1% minor allele frequency. The HGDP dataset is the complete Stanford HGDP SNP Genotyping data. We filtered individuals by removing those not in the “H952” set [13], which leaves individuals without first or second degree relatives in the data. After filtering, the dimensions are 642,951 SNPs by 940 individuals, and a total of 603 million observations (0.08% missing data). The TGP data set was 2012-01-31 Omni Platform Genotypes and is accessible from the NCBI ftp site. We removed related individuals using the sample information provided by the 1000 Genomes Project. After filtering, the dimensions are 1,854,622 SNPs by 1,718 individuals, and a total of 3.1 billion observations (0.3% missing data).

**Simulated data sets.** The goal of our study on synthetic data sets is to demonstrate scalability to tera-sized data sets—one million observed genotypes from one million individuals—while maintaining high accuracy in recovering ground truth per-individual population proportions *θ*_*i*_ and per-population allele frequencies *β*_*k*_. To this end, we generated synthetic genotype data using the Pritchard-Stephens-Donnelly (PSD) model [1]. A specification of the the per-individual population proportions and the population allele frequencies is our “ground truth”. To generate realistic synthetic data, we made the individual *θ*_*i*_’s visually similar to the proportions obtained from the PSD fit to the TGP data set. Further, we modeled allele frequencies *β*_1:*K,l*_ from the same fit.

In Scenario A, the process of drawing an individual *i*’s proportions *θ*_*i*_ has two levels. At the first level, we drew *S* points in the *K*-simplex from a symmetric Dirichlet distribution, *q*_*s*_ *∼* Dirichlet(*α*). Each of the *S* points represents a “region” of individuals, and each individual was assigned to one of the regions such that the regions are equally sized. Then, we drew the population proportions of each individual, *θ*_*i*_ *∼* Dirichlet(*γq*_*s*,1_, …, *γq*_*s,K*_). Thus, each region has a fixed *q*_*s*_ and the proportion of individuals from that region are governed by the same scaled *q*_*s*_ parameter. The parameter *q*_*s*_ controls the sparsity of the *θ*_*i*_, while the parameter *γ* controls how similar admixture proportions are within each group. We set the number of regions *S* = 50, *K* = 6, the Dirichlet parameter *α*= 0.2, and the second-level Dirichlet scale *γ*= 50.

Each *β*_1:*K,l*_ at a SNP location *l*, consists of *K* independent draws from a Beta distribution with parameters following that of the Balding-Nichols Model [14], i.e. 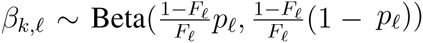 where *p*_*l*_ is the marginal allele frequency and *F*_*l*_ is the Wright’s *F*_*ST*_ at location *l*. The paired parameters *p*_*l*_ and *F*_*l*_ were estimated from the HGDP data set described earlier. For each pair, we chose a random complete SNP from the HGDP data and set the allele frequency *p*_*l*_ to the observed frequency. The Wright’s *F*_*ST*_ *F*_*l*_ was set to the Weir & Cockerham F_ST_ estimate [15] with 5 discrete subpopulations, following analysis of the HGDP study in [16]. We simulated data with 1,000,000 SNPs and three different scales of individuals: 10,000, 100,000 and 1,000,000. With 1 million individuals and 1 million SNPS, the number of observations is tera-sample-sized, i.e., 10^12^ observations.

In Scenario B, each ancestral population is placed at a location evenly spaced along a line (see Figure 3). Individuals are also positioned evenly on the line, and their proportions *θ*_*i*_ are a function of their proximity to each population’s location. This is done by setting a Gaussian density for each ancestral population centered at its location and normalizing each individual such that all proportions sum to 1. For example, choosing *K* = 10 and *N* = 10, 000, we can place the “origin” of each ancestral population at the points *x*_1_ = 1, *x*_2_ = 2, *…, x*_10_ = 10 and place individuals evenly on the interval *x*_*n*_ ∈ [0, 11], e.g. at the points 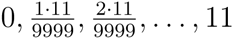. Each individual’s admixture proportions are drawn by first computing each ancestral populations contribution: *f*(*x*_*n*_, *x*_*k*_, *s*), where *f*(*x*, *μ*, *σ*) is the Gaussian density, *x*_*n*_ is the location of individual, *x*_*k*_ (the origin of the *k*th ancestral population) is taken as the mean, and *s* is constant for all populations, taken to be *s* = 2 in our simulations. These contributions are normalized such that they sum to 1. The *β*‘s are generated as in the first scenario.

## The PSD Model

**Modeling assumptions.** We present the model and algorithm for unphased genotype data, though it easily generalizes to phased data. (Most massive population genetics data sets are unphased.) In unphased data, each observation *x*_*i,l*_ ∈ {0, 1, 2} denotes the observed genotype for individual *i* at SNP location *l*. The data are coded for how many major alleles are present: *x*_*il*_ = 0 indicates two minor alleles; *x*_*i,l*_ = 2 indicates two major alleles; and *x*_*i,l*_ = 1 indicates one major and one minor allele. In this last case we do not code which allele came from the mother and which from the father.

The PSD model captures the heterogenous patterns of ancestral populations that are inherent in observed human genomes. It posits *K* ancestral populations, each characterized by its allele frequencies across sites, and assumes that each person’s genome exhibits these populations with different proportions. Given a set of observed genomes, the goal of the algorithm is to estimate (i) the proportion of each ancestral population present in a given individual, (ii) the ancestral population allele frequencies for each SNP, (iii) the effective allele frequency for each individual/SNP combination. Given observed data, we uncover its population structure by estimating the conditional distribution of the allele frequencies and the per-individual population proportions.

Formally, each population *k* is characterized by an array of per-location distributions over major and minor alleles *β*_*k,l*_ ∈ (0, 1). Each individual *i* is characterized by its per-population proportions *θ*_*i,k*_ > 0, where *∑*_*j*_ *θ*_*i,j*_ = 1. The observation for individual *i* at location *l* is assumed drawn from a binomial. Its parameter is a mixture of the population parameters for that location *β*_1:*K,l*_, where the mixture proportions are defined by the individual *θ*_*i*_. Thus, across individuals, the basic population distributions are shared at each location but they are exhibited with different individualized proportions.

Placing priors on the hidden variables, the data are assumed drawn from the following model:

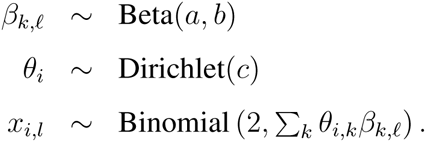

This is the model for unphased data in [1].

**TeraStructure: Scalable Computation for the PSD Model**. How do we use the PSD model? We are given a set of measured genotypes from *N* individuals at *L* locations *x* = *x*_1:*N,*1:*L*_. Given this data, we compute the posterior distribution of the basic population parameters *β = β_1:K,1:L_* and individual population proportions *θ* = *θ* _1:*N,*1:*K*_. From the posterior we can compute estimates of the latent population structure.

For example, Figure S2 illustrates the posterior expected population proportions, computed from our algorithm, for the 1718 individuals of the 1000-Genomes data set. Figure S2 illustrates these posterior estimates at three values of the latent number of populations *K*, at *K* = 7, *K* = 8 and *K* = 9. This data set contains over 3 billion observations. Though the model is not aware of the country-of-origin for each individual, our algorithm uncovered population structure consistent with the major geographical regions. Some of the groups of individuals identify a specific region (e.g., red for Africa) while others represent admixture between regions (e.g., green for Europeans and Central/South Americans).

Specifically, we develop a stochastic variational inference algorithm [7] for the PSD model, a strategy whose computational structure is intrinsically efficient. At each iteration, we first subsample a set of observed genotypes from the data set, a step which involves sampling a location and including the observations for all individuals at that location. We then analyze only those observations at the subsampled location. Finally, we update our estimates of the population-wide hidden structure based on the analysis of the subsample. In each iteration we obtain a new subsample corresponding to a new location and repeat the process.

This is in contrast to previous algorithms for approximate inference in the PSD model, like the MCMC algorithm of [1] or the variational inference algorithm of [3]. These algorithms form an approximate posterior through repeated iterations over the entire data set; such methods are slow for massive data sets. Our method subsamples a SNP location at each iteration, and provides a valid approximation of the admixture posterior that scales to population-size genomic data.

The full algorithm is given in Figure S1. Before deriving its details, however, we describe how we studied and evaluated its performance.

### Experimental setup for the study

The goal of our empirical study is to assess the accuracy and scalability of the stochastic variational inference algorithm of Figure S1 and compare to leading scalable methods in the research literature. In this section, we present the details of our experimental setup. We refer the reader to the main article for the results.

We compared our algorithm to the best existing algorithms for discovering population structure: the fastSTRUCTURE algorithm [3] and the ADMIXTURE algorithm [2]. We fit these algorithms to the largest real-world genotyping data publicly available—the HGDP [8; 9] and the TGP [10] data sets. We also studied fits to massive synthetic data sets. Our synthetic data sets have up to *N* = 1, 000, 000 individuals and *L* = 1, 000, 000 SNP locations, for a total of 10^12^ genotype observations.

On the synthetic data sets, we studied the accuracy of these algorithms in retrieving the ground truth population structure, the run time of these algorithms, and the ability of these algorithms to scale to massive data sets. On the real data sets, we used the predictive approach to evaluating model fitness [12].

**Metrics.** On real data sets, we computed the predictive accuracy on a *test set* of observed genotypes by computing the held-out log likelihood under the PSD model. The test set is chosen to enable a fair comparison to other algorithms. We hold out genotypes for 0.5% of the *N* individuals from each location *l* ∈ 1, … *L*. A better predictive accuracy corresponds to a better fit to the data [12]. We approximate the predictive distribution of a heldout SNP using variational posterior estimates of *θ* and *β*.

On synthetic data sets, we measured the accuracy in recovering the ground truth population proportions. We computed the Kullback Leibler divergence [17] of the variational posterior estimate 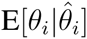 to the true population proportions 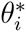 for each individual *i*. We then compared the median KL divergence across all individuals.

**Choosing the number of ancestral populations**. In our experiments on the real data sets (see Table S1), we fixed the number of populations *K* to the optimal values based on validation log likelihoods. Our sensitivity analysis (see Figure S3) revealed that *K* = 8 had the optimal validation likelihood on the TGP data, while *K* = 10 was the optimal for the HGDP data set. In our experiments on simulated data sets (see Table S2), we set *K* to the number of ground truth ancestral populations: *K* = 6.

**Open-source software.** Our software is implemented in C++ and has 5400 lines of code. It uses the POSIX Threading library for multi-threaded computation. It inputs genotype data in text or PLINK format [18] and outputs the population proportions 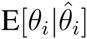. An option to the software tool computes the expected allele frequency Beta parameters local to a list of locations, given the global individual population proportions 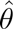 and a list of SNP locations. Our software is available at *http://github.com/premgopalan/terastructure*.

**Computing hardware.** All experiments were run on a single multicore machine with two Intel Xeon E5-2680v2 processors, with 10 cores each and running at 2.8 GHz. The maximum RAM required for our experiments is 10 GB.

### Stochastic Variational Inference for the PSD Model

We now derive the details of our algorithm. We begin by developing a traditional variational inference algorithm for the PSD model. We then extend it to the TeraStructure, the scalable variant. TeraStructure is built on the traditional algorithm.

**Variational Inference for the PSD Model.** The admixture posterior is proportional to the joint distribution

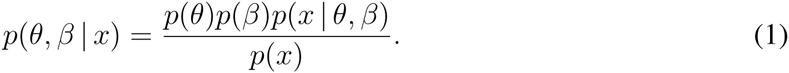

This distribution is difficult to compute because of the normalizing constant, the marginal probability of the observed genotypes. The central computational problem for the PSD model is how to approximate the posterior.

Variational inference is a class of methods for approximate posterior inference that adapts earlier ideas in statistical physics [19] to probabilistic models [5; 6]. Broadly, we define a family of distributions over the hidden variables *q*(*·*) indexed by a set of free parameters *ν*. We then fit *ν* to find the member of the family that is close to the posterior, where closeness is measured with KL divergence,

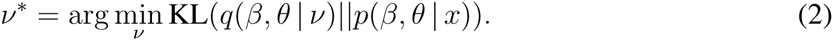

The objective function of Equation 2 is not computable. (It is not computable for the same reason that exact Bayesian inference is intractable—it requires computing the marginal probability of the data.) Thus variational inference optimizes an alternative objective that is equal to the negative KL up to an unknown additive constant,

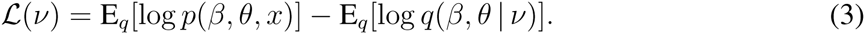

This objective is a function of the variational parameters *ν* because each term is an expectation with respect to *q*(*·*). Further, though the additive constant is unknown, maximizing Equation 3 with respect to *ν* is equivalent to minimizing the KL divergence in Equation 2. Intuitively, the first term encourages that *q*(*·*) place mass on configurations of the latent variables that best explain the data; the second term, which is the entropy of the variational distribution, encourages that *q*(*·*) be diffuse.

To finish specifying the objective, we must set the form of *q*(*·*). A key idea behind variational inference is that the form of the variational distribution is set to make the problem tractable, that is, for the objective of Equation 2 to be computable (as well as its gradients). As for most applications of variational inference, we choose *q*(*·*) to be the *mean-field family*, the family where each variable is independent and governed by its own parametric distribution,

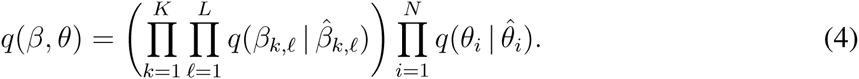

Our notation is that 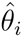 is the variational parameter for the *i*th individual’s population proportions *θ*_*i*_ and 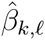 is the variational parameter for the distribution of genotypes in population *k* at location *l*. Further, we set the form of each factor to be the same form as the prior. Thus 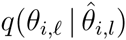 Dirichlet distributions and 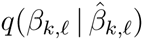 are Beta distributions. These decisions come from the general theory around mean-field variational inference in exponential families [20; 21]. (See also Equation 7 and Equation 9.)

We emphasize that in the variational family each hidden variable is endowed with its own variational distribution. While the model assumes each individual’s proportions come from the same shared prior, the variational family provides a different parameter for each. This gives the variational family the flexibility it needs to represent different individuals with different population proportions. For example, to create Figure S2 we plotted the variational expectation of each individuals population parameters distribution 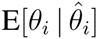.

With these components—the objective of Equation 3 and the variational family of Equation 4—we have turned the inference problem for the PSD model into an optimization problem.

**Stochastic Variational Inference for the PSD Model.** Traditional variational inference iterates over all the variational parameters. For example, the authors of [3] approximate the admixture posterior by updating each variational parameter in turn while holding the others fixed. This *batch* strategy is more efficient than MCMC but cannot scale to tera-sized data sets, where the number of individuals *N* is in the hundreds of thousands or millions and the number of locations *L* is in the millions.

To solve this problem, we use stochastic optimization [22] applied to the variational objective [23; 7]. Our algorithm follows easy-to-compute noisy estimates of the gradient to more quickly make progress in the variational objective. The noise and computability of the gradient stem from repeated subsampling from the data.

TeraStructureaintains variational estimate of each individuals population proportions 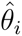 and the allele frequencies of each basic population 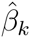. It repeatedly cycles through the following steps:

1. Sample an observation from SNP location *l* from all individuals.
2. Estimate how each individual expresses the basic populations, only using the measured location *l*. Use these estimates to update the ancestral allele frequencies parameter 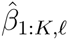 local to *l*.
3. Use these estimates to update the population proportions parameter of all individuals, 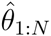.

The stochastic algorithm above can quickly make progress. After one iteration, which involved processing observations at only one of the *L* possible locations, we have an estimate of the population proportions of all individuals. Given the estimate of the population proportions, an estimate of the ancestral allele frequencies can be computed for any location. In comparison, the batch algorithm of [3] needs to iterate over the entire data set at least once, to make any progress.

**Global and local parameters.** Before we develop our algorithm, we use the conditional dependencies in the model to divide the variational parameters into *local* and *global* parameters [7].

In each iteration we subsample genotype measurements for all individuals at a SNP location *l*. Our sampled observations are *x*_1:*N,l*_. Under the PSD model, given individual proportions *θ*_1:*N*_, the sample *x*_1:*N,l*_ and the ancestral allele frequencies β_1:*K,l*_ are conditionally independent of all other observations and allele frequencies *β*_1:*K,-l*_. Thus, the allele frequencies β_1:*K,l*_ are local to the observations *x*_1:*N,l*_. The population proportions *θ*_1:*N*_, with the local variables, govern the distribution of observations at any sampled SNP location. Therefore, the *θ*_1:*N*_ are global variables. Following [7], we extend this notion of global and local sets to the variational parameters. Given observations *x*_*l,*1:*N*_ at the location *l*, the 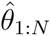 are the global variational parameters; the 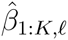 are the local variational parameters.

In stochastic variational inference [7], we iteratively update local and global parameters. In each iteration, we first subsample a SNP location *l* and compute optimal local parameters for the sample, given the current settings of the global parameters. We then update the global parameters using a stochastic natural gradient [24] of the variational objective computed from the subsampled data and the local parameters.

We will now develop our algorithm by first obtaining closed form updates for our local and global variational parameters. For the local parameters, we will derive optimal coordinate updates; for the global parameters, we will derive the stochastic natural gradient update.

**Computing the optimal local parameters.** Given the global parameters 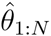, we can optimize local parameters 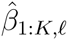 in closed form under certain assumptions. These assumptions involve the *complete conditionals* of the hidden variables in the model, and the variational family. A complete conditional is the conditional distribution of a latent variable given the observations and the other latent variables in the model [20]. If the complete conditional of a variable is in the same family as its prior, and the corresponding variational distribution is in the same family, then we can optimize its variational parameter by setting it to the expected natural parameter (under *q*) of the complete conditional.

If the complete conditional of each latent variable is in the same exponential family as its prior distribution, then the model is *conditionally conjugate*.

The complete conditionals for the *β*_*k,l*_ at a sampled location *l* are

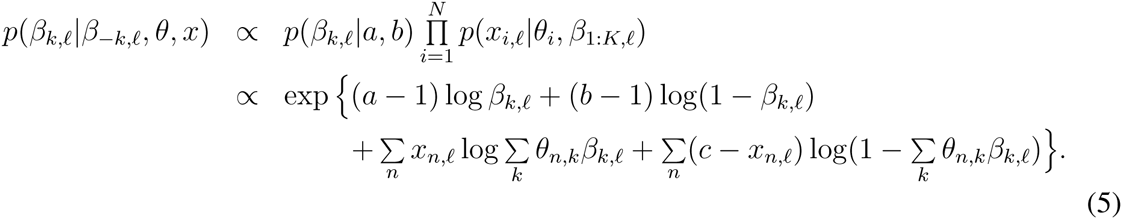

The complete conditional in Equation 5 is not in the exponential family because the expectation of the second and third log-of-summation terms, with respect to the variational family *q*, are intractable. Therefore, the PSD model is not conditionally conjugate.

To overcome the nonconjugacy in the model, we introduce multinomial approximations using the zeroth order delta method for moments [25; 26]. These approximations provide a lower bound to these intractable terms in the variational objective of Equation 3. In particular, we introduce auxiliary *K*-multinomial distributions *q*(*ϕ*_*il*_) and *q*(*ξ*_*il*_),

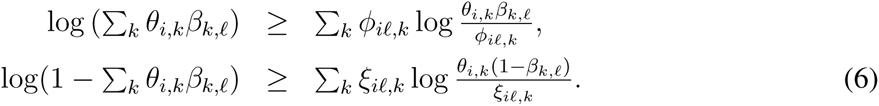

These distributions *q*(*ϕ*_*il*_) and *q*(*ξ*_*il*_) approximate only the conditionals of the allele frequencies local to the sampled location *l* and the individual *i*; the parameters to these distributions are local.

Substituting the lower bounds from Equation 6 in Equation 5, the complete conditional is

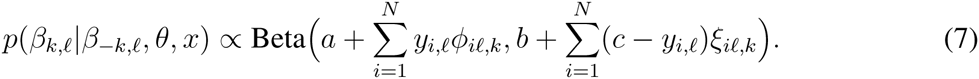

Our approximation has effectively placed the complete conditional of allele frequency *β*_*k,l*_ in the exponential family. By choosing the variational distribution 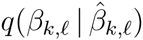 from Equation 4 to be the Beta distribution, the same family as the prior distribution, we satisfy the conditions for a closed form coordinate update for the local parameters 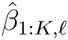. The optimal 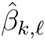 is the expected natural parameter (under *q*) of the complete conditional in Equation 7 [20].

Another perspective on the approximations in Equation 6 is they lead to a computationally efficient lower bound on the objective of Equation 3.

**Computing stochastic gradient updates for the global parameters**. We now turn to the stochastic optimization of the population proportions parameter 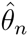 using the subsampled observations *x*_1:*N,l*_ at location *l*. We compute noisy estimates of the natural gradient [24] of the variational objective with respect to 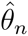, and we follow these estimates with a decreasing step-size. Following [7], we can compute the natural gradient of Equation 3 with respect to the global variational parameter 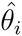 by first computing the coordinate update for 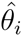 and then subtracting its current setting. To compute the coordinate update for 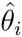, we write down the complete conditional of the population proportions *θ*_*i*_:

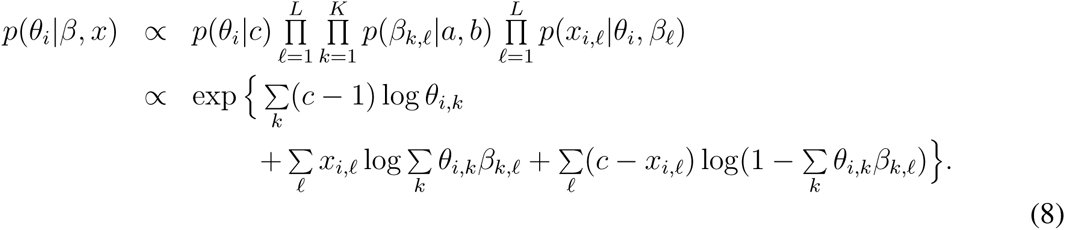

Similar to the complete conditionals of the local variables in Equation 5, the complete conditional in Equation 8 is not in the exponential family. We use the multinomial approximations in Equation 6 to bring the complete conditional into the exponential family, and in the same family as the prior distribution over the population proportions:

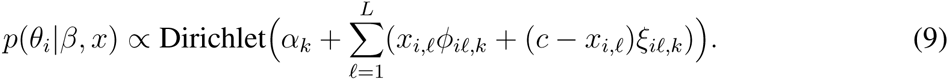

Following [7], the stochastic natural gradient of the variational objective with respect to the global parameter 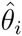, using *L* replicates of *x*_*i,l*_ is

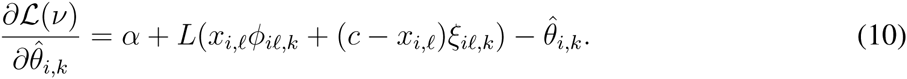

Notice we have used the expected natural parameter from the complete conditional in Equation 9 in Equation 10. We arrive at this form of the natural gradient by premultiplying the gradient by the inverse Fisher information, and replacing the summation over all SNP locations in Equation 10 with a summation over *L* replications from the sampled location. Equation 10 is a noisy natural gradient of a lower bound on the variational objective of Equation 3.

To optimize the variational objective with respect to the population proportions 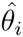, we use the natural gradients in Equation 10 in a Robbins-Monro algorithm [7]. At each iteration we update the global variational parameters with a noisy gradient computed from the SNP observations at location *l*. The step-size at iteration *t* is *ρ*_*t*_, and is set using the schedule

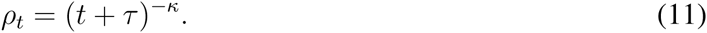

This satisfies the Robbins-Monro conditions on the step-size, and guarantees convergence to a local optimum of the variational objective [22].

**The stochastic algorithm.** The full algorithm is shown in Figure S1. For each iteration, we first subsample a SNP location *l* and compute optimal local parameters 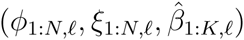 for the sample, given the current settings of the global parameters 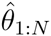. We then update the global parameter 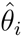 of each individual *i* using the stochastic natural gradient of the variational objective, with respect to 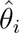, computed from the subsampled data and local parameters.

**Memory efficient computation.** During training, the stochastic variational inference algorithm is only required to keep the variational population proportions 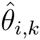 for all individuals *i* ∈ 1, …, *N* in memory. For a given location, the optimal local parameters 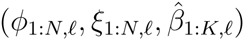 can be computed using the local optimization steps—steps 6 to 9—in Figure S1. The local parameters need not be kept around beyond the corresponding sampling step. This drastically cuts the memory needed. The memory requirement is therefore *O*(*NK*) where *N* is the number of individuals and *K* is the number of latent ancestral populations. Further, this results in a small fitted model state: the fitted 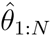. Given the 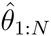, the allele frequencies 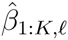 can be optimized for any given location *l*, using the local step.

**Linear scaling in the number of threads.** We can compute the local steps and the global steps in parallel across *T* threads. First, we “map” the individuals into *T* disjoint sets, and each thread is responsible for computation on one of these sets of individuals. Notice that each thread can independently compute the local parameters (*ϕ*_*n,l*_, *ξ*_*n,l*_) for any individual *n* that it owns. This corresponds to step 6 of the algorithm in Figure S1. Further, the sums required in step 7 of the algorithm in Figure S1 can also be computed in parallel. The “reduce” step consists of aggregating the per-thread sums in step 7, and estimating the new Beta parameters. This is an *O*(*T* + *K*) operation, where *T* is the number of threads and *K* is the number of ancestral populations. Since *T* and *K* are small constants, our reduce step is inexpensive. The global step in step 9 can also be computed in parallel.

Given *T* threads, the computational complexity of the stochastic algorithm is 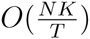. The algorithm is dominated by the parallel computation in steps 6 and 9, which scale linearly in the number of threads *T*. By increasing *T*, we scale our algorithm linearly in the number of threads.

**Initializing variational parameters.** We initialize the population proportions randomly using *θ*_*ik*_ ∼ Gamma(100, 0.01). Within each local step, we initialize 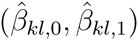 at location *l* to the prior parameters (*a, b*). We use the same initialization procedure on all data sets.

**Assessing convergence using a validation set.** We hold out a *validation set* of genotype observations, and evaluate the predictive accuracy on that set to assess convergence of the stochastic algorithm in Figure S1 [12]. (These observations are treated as missing during training.)

The validation set is chosen with computational efficiency in mind. We will periodically evaluate the heldout log likelihood on this set (the *validation log likelihood*) to determine convergence of the algorithm in Figure S1. By choosing individuals from a small fraction of total locations *L*, we ensure that this periodic computation is only required to recompute the optimal 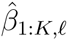 for those locations.

The TeraStructure algorithm stops when the change in validation log likelihood is less than 0.0001%. We measure this change over 100, 000 iterations.

For the validation set, we uniformly sample at random 0.5% of the *L* locations, and at each location we uniformly sample at random and keep aside observed genotypes for *r* individuals. The number of per-location held out individuals *r* is set to *N/*100 for large *N* (*N* > 2000) and otherwise to *N/*10. This allows for a reasonably small fraction of individuals to be held out from each location. Further, *r* is limited to a maximum of 1000 individuals for any *N*.

**Hyperparameters.** We set the Dirichlet parameter *c* to 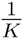 to enforce a sparse prior on the per-individual population proportions. We set the learning rate parameters, *τ*_0_ to 1 and *κ* to 0.5, to allow rapid learning in the early iterations. Finally, we set the hyperparameters *a* and *b* to 1 to enforce a uniform prior on the population parameters *β*_1:*K,*1:*L*_. We used the same hyperparameter settings and initialization in all of our experiments.

## Supplementary Figures and Tables

**Figure S1:**
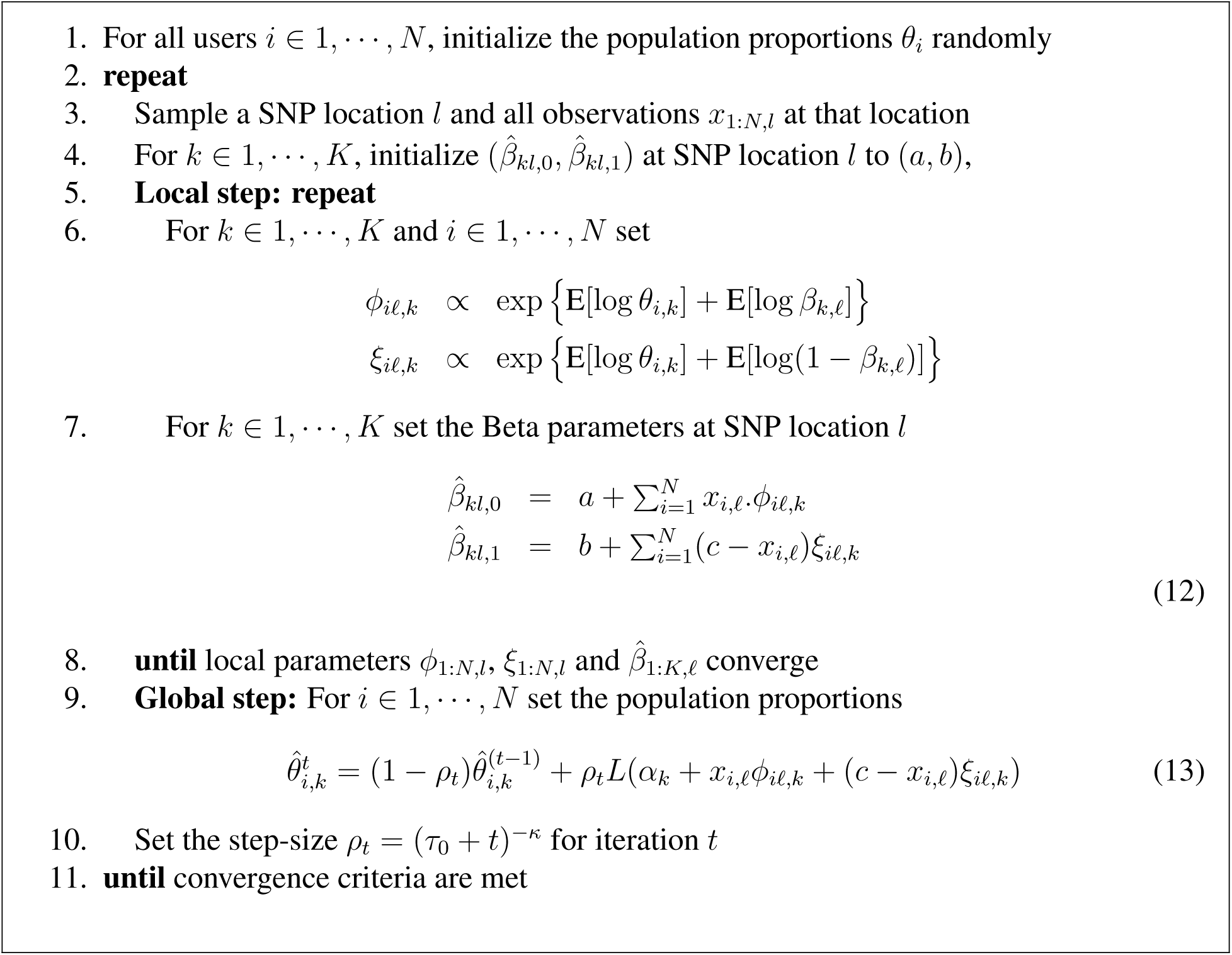
TeraStructure Algorithm – Stochastic variational inference for the PSD model.

**Figure S2:**
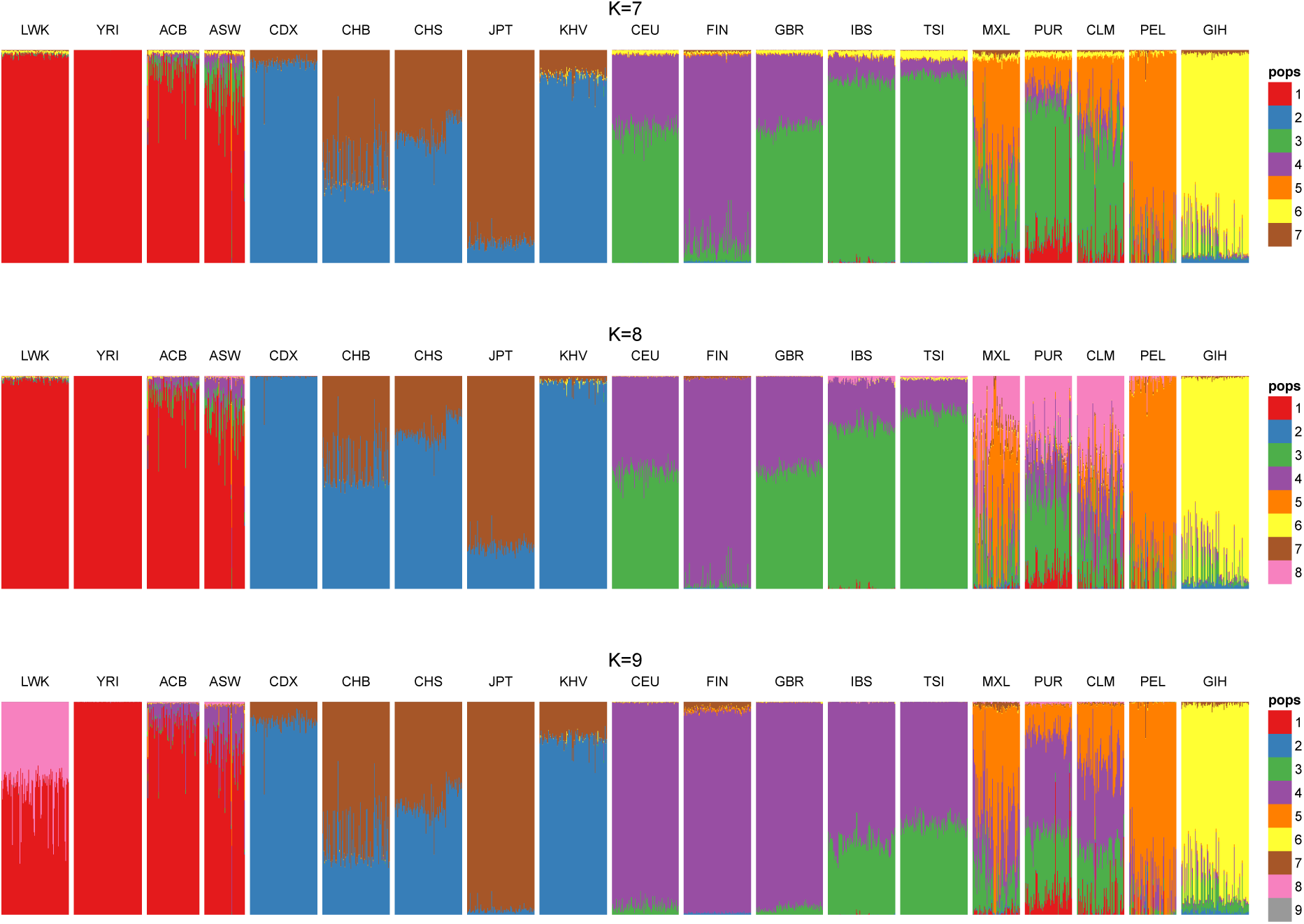
Population structure inferred from the TGP data set using the TeraStructure algorithm at three settings for the number of populations *K*. The visualization of the θ‘s in the Figure shows patterns consistent with the major geographical regions. Some of the clusters identify a specific region (e.g. red for Africa) while others represent admixture between regions (e.g. green for Europeans and Central/South Americans). The presence of clusters that are shared between different regions demonstrates the more continuous nature of the structure. The new cluster from *K* = 7 to *K* = 8 matches structure differentiating between American groups. For *K* = 9, the new cluster is unpopulated.

**Figure S3:**
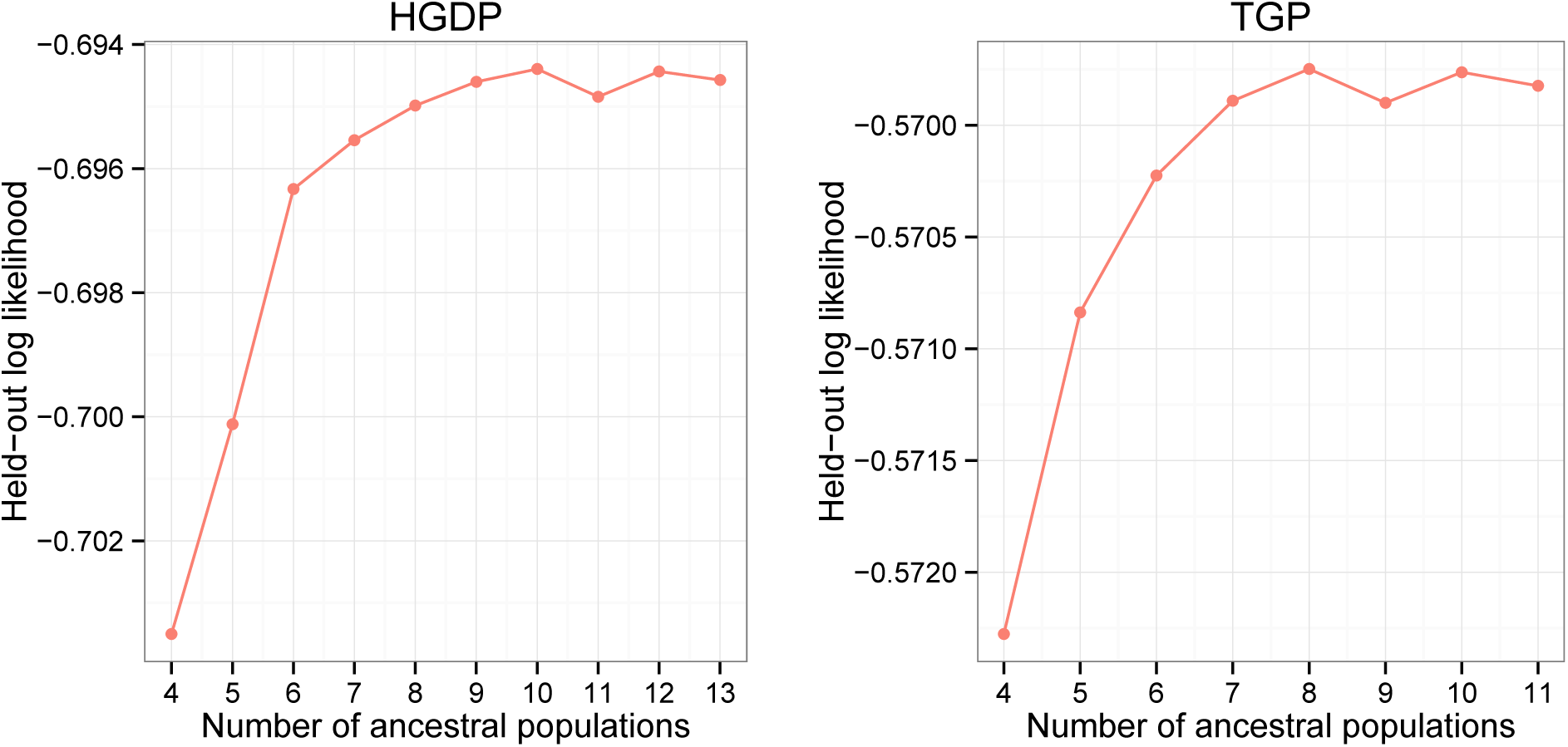
Predictive log likelihood as a function of the number of ancestral populations on the Human Genome Diversity Panel (HGDP) and 1000 Genomes Project (TGP) data sets. The HGDP data peaks at 10 population, and the TGP data peaks at 8 populations.

**Table S1:** The predictive accuracy of TeraStructure is comparable to the ADMIXTURE [2] and the fastSTRUCTURE [3] algorithms, implying a similar model fit. The mean test log likelihood under the model fits is shown. We generated 5 test sets at random and computed the mean over these heldout sets. *N* is the number of individuals in the data set. The number of ancestral populations is set to *K* = 10 for HGDP and *K* = 8 for TGP.

| Data set | N | Mean predictive log likelihood |  |  |
| --- | --- | --- | --- | --- |
|  |  | TeraStructure | ADMIXTURE | fastSTRUCTURE |
| HGDP | 940 | -0.71 | -0.71 | -0.71 |
| TGP | 1718 | -0.60 | -0.60 | -0.61 |

**Table S2:** The accuracy of the algorithms on synthetic data generated via Scenario A. TeraStructure is the only algorithm that was able to complete its analysis on the synthetic data sets with *N* = 100, 000 individuals and *N* = 1, 000, 000 individuals. On these massive data sets, TeraStructure found a highly accurate fit to the data (see also Figure 2). On smaller synthetic data, TeraStructure finds a fit to the data that is closer to the ground truth than either of the other methods. The number of ancestral populations is set to the number of ground truth ancestral populations: *K*=6.

| Data set | Replication | $N$ | $L$ | Median per-individual KL divergence | | |
| --- | --- | --- | --- | --- | --- | --- |
|  |  |  |  | TeraStructure | ADMIXTURE | fastSTRUCTURE |
| Scenario A (10K) | 1 | 10,000 | 1,000,000 | **0.016** | 0.020 | 6.68 |
| Scenario A (10K) | 2 | 10,000 | 1,000,000 | **0.009** | 0.019 | 5.15 |
| Scenario A (10K) | 3 | 10,000 | 1,000,000 | **0.020** | 0.022 | 4.49 |
| Scenario A (100K) | 1 | 100,000 | 1,000,000 | **0.006** | – | – |
| Scenario A (100K) | 2 | 100,000 | 1,000,000 | **0.013** | – | – |
| Scenario A (100K) | 3 | 100,000 | 1,000,000 | **0.009** | – | – |
| Scenario A (1M) | 1 | 1,000,000 | 1,000,000 | **0.015** | – | – |
| Scenario B | 1 | 10,000 | 100,000 | **0.21** | 0.42 | 7.21 |
| Scenario B | 2 | 10,000 | 100,000 | **0.27** | 0.42 | 7.97 |
| Scenario B | 3 | 10,000 | 100,000 | **0.26** | 0.42 | 7.68 |
| Scenario B | 1 | 10,000 | 1,000,000 | **0.16** | – | – |
| Scenario B | 2 | 10,000 | 1,000,000 | **0.23** | – | – |
| Scenario B | 3 | 10,000 | 1,000,000 | **0.25** | – | – |

## Footnotes

1 We describe and illustrate these quantities as though they are estimates. More technically, the algorithm stores parameterized distributions of them.

